# Towards a standardised method for the characterisation and isolation of colorectal cancer stem cells by SdFFF and UHF-DEP: highlighted by transcriptomic analysis

**DOI:** 10.1101/2024.11.04.621916

**Authors:** Charlotte Jemfer, Nina Blasco, Stéphanie Durand, Elisa Blin, Robin Mas, Isabelle Basly, Marie Boutaud, Héloïse Daverat, Arnaud Pothier, Claire Dalmay, Muriel Mathonnet, Gaelle Begaud, Serge Battu

## Abstract

Cancer stem cells (CSCs) play a crucial role in tumor heterogeneity and the progression of colorectal cancer (CRC). However, identifying and isolating CSCs remains challenging, as conventional methods, such as fluorescent or magnetic labeling, are time-consuming and require substantial technical and biological resources, limiting their clinical applicability. In this study, we evaluated the potential application of a previously established approach with glioma cells, combining sedimentation field-flow fractionation (SdFFF) with a label-free Ultra-High Frequency dielectrophoresis (UHF-DEP) biosensor, for isolating and identifying CRC CSC. The integration of the UHF-DEP biosensor with SdFFF enables the identification of CSCs without specific labeling, simplifying their analysis. To optimize this coupling, we standardized the mobile phase between the two technologies. Producing two CSC sub-populations eluted in Fraction 1 and Fraction 3. For the first time, transcriptomic analysis was employed to deepen our understanding of the genetic composition of the cells, complementing functional and phenotypic characterizations. From these characteristics, it emerged that F1 cells corresponded to precursors, while F3 cells could be considered CSC. The UHF-DEP biosensor demonstrated its ability to detect these differences, linking F3 to cells cultivated in defined medium (DM), used as a reference standard. Finally, transcriptomic analysis (RNA-seq) has deepened our understanding of the genetic profiles of these subpopulations, enriching the interpretation of their characteristics as CSCs. This in-depth approach will also contribute to a better understanding of the results obtained by the UHF-DEP biosensor.

## 1. Introduction

Colorectal cancer (CRC) is the third deadliest cancer worldwide [1]. Despite standard treatment, which combines surgical resection and adjuvant chemotherapy, around 50% of CRC patients experience relapse within two years of initial treatment [2]. This phenomenon is largely attributed to the therapeutic resistance of colorectal cancer stem cells (CSCs) [3]. CSCs play a pivotal role in tumor development, disease progression, and recurrence [4]. Due to theirs multipotency and self-renewal abilities, these cells contribute to the tumor growth. They supply a reservoir of undifferentiated cells able to generate new tumor cells which sustain the tumor mass [5].

Furthermore, CSCs are involved in metastatic spread, as they can revert to an epithelial phenotype, facilitating migration and colonization of distant organs, contributing to poor prognosis in CRC [6]. A major challenge in treating CSCs is their quiescence, a dormant state that allows them to evade standard treatments like chemotherapy, which primarily target actively dividing cells [7]. This ability to remain inactive and resist therapy often leads to relapse, as surviving CSCs can later reactivate, driving the regrowth of the tumor after an initial response to treatment. This contributes to the phenomenon of minimal residual disease (MRD), where a small number of CSCs persist undetected after treatment, leading to the eventual recurrence of the cancer [6]. CSCs also contribute to tumor heterogeneity, complicating therapeutic strategies due to the presence of different subpopulations of cells that may respond variably to treatment. Adding to this complexity is their remarkable cellular plasticity—the ability of CSCs to shift between stem-like and more differentiated states depending on environmental cues and treatment pressures [8]. This plasticity enhances their capacity to survive in various microenvironments, further complicating their targeting.

At present, CSCs are typically identified using a combination of phenotypic markers and functional assays. Expression or overexpression of phenotypic markers, explored by FACS or MACS, include the analysis of surface proteins such as *CD133, EPCAM, CD44* and so forth, as well as intracellular markers like *NANOG, OCT4, LGR5*, or *SOX2* [5], [9]. Functional assays, such as ALDH activity measurement [10], clonogenic assays, and *in vivo* xenograft models (gold standard) [9], [11], are used to complet phenotypic identification and confirm CSC characteristics. However, due to the cellular plasticity of CSC, marker expression can fluctuate, making it necessary to combine both phenotypic and functional characterizations for accurate identification. This dual approach is crucial, as relying on phenotypic markers alone may overlook certain CSC subpopulations or lead to uncertain identifications due to marker variability.

Then the combining use of these methods make diagnostics long and expensive process. Therefore, it is essential to develop a tool capable of quickly providing and identifying an enriched CSC population from a heterogeneous cancer cell population, without inducing any alterations [12]. Based on previous study [13], the hyphenation of two label-free techniques have emerged as promising alternatives for CSC isolation and characterization: sedimentation field-flow fractionation (SdFFF) as cell sorting method and ultra-high frequency dielectrophoresis (UHF-DEP) as characterization one.

These techniques preserve the functional integrity of the cells, avoiding the drawbacks of labelling such as differentiation or cellular degradation. SdFFF enables the isolation of CSCs based on their biophysical properties, including size, density, and rigidity, by applying hydrodynamic lift forces and a perpendicular multigravitationnal external field in a ribbon-like separation channel. The cells are separated under the “Hyperlayer” elution mode, driven by size and density, in a gentle and non-invasive process [14], [15], [16]. On the other hand, the UHF-DEP biosensor characterizes cells based on their cytoplasmic dielectric properties by measuring their electrical response to an ultra-high frequency (50-600 MHz) electric field, enabling the detection of subtle differences in cytoplasmic and nucleus composition without causing damage to the cells [13], [17]. UHF-DEP characterization offers a rapid, precise, and cost-effective way to distinguish between subpopulations of CSCs and differentiated cells based on their dielectric traits.

The combination of SdFFF and UHF-DEP has already been successfully applied to glioblastoma [13], where it enabled the isolation and characterization of glioblastoma CSCs. In this study, we aim to explore whether this coupled approach can be extended to other solid tumors, specifically colorectal cancer (CRC). Our goal is to validate the use of this system for isolating and characterizing CRC CSCs, adapting the experimental conditions to ensure compatibility with the UHF-DEP detector.

A key aspect of this adaptation involves switching the mobile phase in SdFFF to the DEP medium, which is essential for UHF-DEP characterization. The DEP medium, composed of 8.5% sucrose, provides low conductivity, making it compatible with the UHF-DEP process [18]. This medium maintains osmotic balance, ensuring that the cells remain unaltered and viable upon contact. However, it displayed different physicochemical properties compared to the PBS originally used in SdFFF [14], [15], in particular concerning mobile phase density and viscosity, modifying sample behaviour. In consequences and in contrast to the previous results obtained for Glioblastoma [13], elution conditions have to be re-optimized, involving a new and crucial biological calibration of fractograms to ensure the ability of the system to isolate CSCs.

In addition, to enhance this biological characterization, we have extended the identification of each population (F1-F3-PT) for the first time to RNA-Seq analysis. This initial identification of fractions by transcriptomics will enable a deeper and more precise understanding of the nature of the cells subjected to characterization by the biosensor, and will also contribute to a better understanding of the intracellular characterization it performs. In addition, a calibration step using a cellular gold standard is also essential, to serve as a reference for the biosensor.

By extending the SdFFF and UHF-DEP hyphenation in CRC diagnosis, we aim to establish a more standardized, label-free method for CSC isolation and characterization. This could pave the way for broader applications across various solid tumors, ultimately contributing to more effective CSC-targeted therapeutic strategies and improved cancer management, particularly in CRC.(SI 1)

## 2. Materials and methods

### 2.1. Cell culture

SW480 cell line were purchased from the American Type Culture Collection (ATCC/LGC Promochem, Molsheim, France) and were grown in RPMI 1640 GlutaMAX™ (Gibco™, Thermo Fisher, MA, USA) medium supplemented with 10% FBS, 1%sodium pyruvate and 1% of penicillin / streptomycin (Thermo Fisher) at 37°C under a humidified atmosphere of 5% CO2.

Prior to each SdFFF sort, cells are seeded at a density of 1×10^6^ cells/mL in two T75 flasks in 10 ml medium for 72h.

To enrich the cell population in CSCs, cell lines or CPP were cultured for at least one week in a “Define Medium” (DM): DMEM/F-12 (Thermo Fisher) medium without FBS but supplemented with 5 µg/mL of insulin, N-2 supplement (1X), 1% of penicillin / streptomycin (Thermo Fisher), 20 ng/mL of EGF and 20 ng/mL of bFGF (MACS Miltenyi Biotec, Bergisch Gladbach, Germany).

### 2.2. SdFFF device and cell elution conditions

#### 2.2.1. SdFFF cell sorting

The SdFFF separation technology used in this study has been widely described [19]. The elution conditions developed for optimum separation of cells in DEP medium are as follows:

|  |  |
| --- | --- |
| SW480 Cell suspension : | $2.5 \times 10^6$ cells/mL |
| Flow rate : | 1,0 mL/min |
| Mobile phase : | Sterile DEP medium, pH 7.4 and conductivity 20 mS/m |
| Multi-gravitational field : | 12 g (270 rpm) |

Each of the fractions was collected at separate times: F1: 3’20 to 4’30 and F3 6’ to 7’50. A third fraction is also collected, constituting the total eluted population (PT) without the void volume, making it possible to constitute an internal control from 3’20 to 7’50. The initial population also constitutes the remaining unsorted cell suspension (CR) and is used as an external control population. In order to obtain a sufficient quantity of cells for characterization analyses, consecutive collections (9-13 injections of 100µl each) were carried out.

#### 2.2.2. Cell size characterization

In this study 4 collections of fractions (F1/F3/CR) and 2 collections of total peak (PT) were made. The Multisizer 4 was used to measure the average cell size of a sample. The cells were then added to 10 ml of Isoton and between 20,000 and 50,000 cells were analysed. The analysis were performed for each biological replicate.

### 2.3. Biological Cell characterizations

#### 2.3.1. RTqPCR

Après avoir réalisé les collections de fractions en sortie de SdFFF RNA extraction and RTqPCR analysis were extracted using the RNeasy Mini Kit (QIAGEN) and quantifed by NanoDrop 2000 (Termo Fischer). For RT-qPCR analyses, the RNA were reverse-transcribed into cDNA using the cDNA Archive kit (Applied Biosystems). Quantitative gene expression was performed using SensiFAST Probe Hi-ROX kit (Bioline, London) on a QuantStudio 5 (Termo Fischer). Results were normalized to GAPDH expression and analyzed using the ΔΔCt method. Every experiment were performed in triplicate from each biological replicate.

| Gene | Supplier | Reference |
| --- | --- | --- |
| GAPDH (GAPDH) | Thermofisher | Hs02758991_g1 |
| NANOG (Nanog) | Thermofisher | Hs02387400_g1 |
| POU5F1 (Oct4) | Thermofisher | Hs00999634_gH |
| SOX2 (Sox2) | Thermofisher | Hs01053049_s1 |
| LGR5 (LGR5) | Thermofisher | Hs00969422_m1 |

#### 2.3.2. Cellular cycle

The cells were centrifuged at 300 RPM for 10 min. A first wash was then carried out by removing the supernatant and recovering the cell pellet in 2 mL of PBS (Thermofisher scientific, France) at 4°C before centrifuging at 1500 RPM for 5 min. Following this first wash, 2 other successive washes were carried out under the same conditions. After the washes, the cells were suspended in 300 μL of PBS at 4°C, then 700 μL of ethanol at -20°C was quickly added by vortexing the tube. Samples were then stored overnight at -20°C. Before analysis, the tubes were placed at RT for 10 minutes, then centrifuged at 1500 RPM for 5 min. The supernatant was removed with a pipette and 2 mL of cold PBS was added. The wash was repeated three successive times (1500 RPM for 5 min at 4°C) until the Ethanol was removed. After washing, the cells were suspended with 400 ml of PBS and 24 μL of RNase A. After 20 min incubation at RT, 50µl of propidium iodide is added to each sample prior to FACs Calibur analysis.

#### 2.3.3. Clonogenicity assay

After SdFFF sorting, 1000 cells from each condition were placed in 100µl of Matrigel matrix ref : 356255 in a 96-well plate and incubated for 15 min. Next, 100 µl of culture medium at 37°C was gently applied to the polymerised matrigel. 50 µl was removed after 7 days and 50 µl of new medium was added to each well. Image monitoring was carried out every week for 3 weeks. Observed at 10x magnification.

The area of the spheroids observed is measured for each condition. Images of the wells were then taken by a fluorescence stereo zoom microscope (LEICA MZFLIII, Wetzlar, Germany), and images were analyzed by ImageJ.

#### 2.3.4. Proliferation

To evaluate proliferation, phase confuence was determined and normalized with t=0 h using the Incucyte analysis software

#### 2.3.5. Proteome profiler

The proteome profiler human pluripotent stem cell array kit (R&D Systems) was used to evaluate the expression of cancer stem cell markers, according to the manufacturer’s instructions. Briefly sorted cell lysate proteins were extracted using RIPA buffer (Sigma-Aldrich) and sonication (VibraCell™). According to the manufacturer’s instructions 10 µg / membrane was used, and the chemiluminescent signal was detected using a Chemidoc system (BioRad). ImageJ software was used to quantify the pixel density according to the Array’s manual.

#### 2.3.6. RNAseq

##### Gene expression profiling by RNA-Seq and bioinformatics analysis

Sorted cells were stored at -20°C in lysis buffer for at least 24 hours, after which mRNA was extracted using the Quick-RNA™ Microprep Kit (Zymo Research). Each sample’s quality and concentration were assessed with a Nanodrop and the Agilent RNA 6000 Nano Kit (Agilent Technologies). Only samples with a concentration of at least 50 ng/µl and an RNA Integrity Number (RIN) greater than 7 were included for further analysis.

RNA sequencing was conducted by the Beijing Genomics Institute, following the manufacturer’s protocol for Rolling-Circle and DNA Nano-Ball (DNB™) technology. Paired-end reads were generated with a length of 150 bases using the DNBSEQ platform, with an average yield of 10.19G data per sample. Bioinformatics analysis was conducting by the Beijing Genomics Institute according following analysis pipeline: 1) Raw data obtained from sequencing were filtered using SOAPnuke[20] to remove reads with low quality (reads with a percentage of bases with a quality value less than 15 exceeding 20% of the total bases in the read), adapter contamination, and high content of unknown bases (N) (greater than 0.1%). More of 30 million high-quality (clean) paired reads per sample were obtained after read filtering step, with more 90% of reads with quality greater than Q30 (range: 89.85 to 96.34%). 2) Clean data was aligned to the reference gene set using Bowtie2 [21](v2.3.4.3) and gene expression quantification was performed using RSEM [22](v1.3.1) software. Alignment results show a range of 78.74 to 84.96% of read uniquely mapped to reference. Finally, the average alignment of the gene set was 87.20% and a total of 18856 genes were detected.

Differential gene expression was conducted pair-wise using DESeq2[23] to identify differentially expressed genes (DEGs) in different cells fractioned by sdFFF (F3 *versus* PT, F1 *versus* PT and F3 *versus* F1) according to the following selection criteria: absolute log_2_ fold change superior to 1 and p-adjusted (after Benjamini-Hochberg (BH) multiple testing correction) inferior to 0.05. To increase detection power of DEGs, filtration of lowly expressed genes was previously performed using the R package HTSFilter[24]. The results of differential analysis were visualized using volcano plot (EnhancedVolcano package [25]) and hierarchical clustering analysis (ComplexHeatmap package)[26]. Principal component analysis was performed using FactoMineR and factoextra packages[27].

To further explore gene functions related to phenotypic changes, we conducted GO (http://www.geneontology.org/) and KEGG (https://www.kegg.jp/) and REACTOME (https://reactome.org/) enrichment analysis on differential expressed genes using DAVID [28], [29] and only retained entries with p-adjusted inferior to 0.05 using the Benjamini-Hochberg correction. Reduction of redundancy through the grouping of biologically similar terms is performed by using “functional annotation clustering” tool implemented in DAVID, that permit to group annotation terms together that exhibit similar associations with DEGs list.

##### Data integration into biological networks

To obtain biological information about set of deregulated genes in F3 as compared to F1 fraction, we searched for direct (physical) and indirect (functionally associated with no direct interaction) associations between proteins coding by differentially expressed genes in the STRING [30](Search Tool for the Retrieval of Interacting Genes/Proteins) database (version 12.0). Briefly, interactions in STRING are derived from multiple sources (experimental/biochemical experiments, curated databases, genomic context prediction, co-expression and automated text mining). Confidence level of edge, i.e. interaction between 2 nodes or protein, is computed in a combined score (ranging from 0 to 1). Interactions and transcriptome data were imported and used for network generation in the Cytoscape [31] environment (version 3.10). Topological analysis was performed with Cytohubba [32]to identify nodes parameters, including degree and bottleneck.

### 2.4. Biosensor operation and characterization of DEP medium

#### 2.4.1. DEP medium

DEP-B consists of 85.56 g sucrose, 0.12 g Tris Base and 0,066 g anhydrous MgCl_2_. Each component is diluted in 800 mL distilled water, with stirring. Once the solution is homogeneous, the pH is adjusted to 7.4 (physiological pH) using concentrated HCl solution. The mixture is then made up to 1 L in a volumetric flask, sterilized by 0.02µ filtration and stored at 4°C for up to one month.

#### 2.4.2. DEP characterization

##### 1.2.1.1. Viscosity

Viscosity measurements are carried out using the Rhéomat RM 200 on 3 DEP-Bs from different production facilities and compared with industrial PBS (Thermofisher scientific, France), at room temperature.

##### 1.2.1.2. Density

To measure the density of DEP-B, volumes ranging from 1 mL to 40 mL were progressively weighed into an empty container. Density is calculated by dividing the mass of the liquid by its volume, expressed in g/mL.

##### 1.2.1.3.Osmolarity

The osmolarity of DEP-B was measured on 3 independently produced samples using the OsmoPRO® Osmometer.

#### 2.4.3. UHF-DEP characterization

In our project, we direct the cells into a micro-fluidic channel consisting of a PDMS (Polydimethylsiloxane) integrated above a BiCMOS sensor (**Erreur ! Source du renvoi introuvable**.A). The sensor consists of several 40 × 40µm quadrupoles, with 4 electrodes placed in the center of the channel (2 thin 0.45 µm thick electrodes placed parallel to the flow and 2 thicker 9 µm electrodes, perpendicular to the flow of cells).

The biosensor consists of a micro-fluidic chip that applied the frequencies at which cell movement is observed. More specifically, a variation of pressure pushes the cells to a quadruple, where they’re first subjected one by one to an ultra-high-frequency (MHz) field at negative DEP forces (**Erreur ! Source du renvoi introuvable**.B). Once stabilized, we progressively reduce the electric field until the cell moves towards one of the quadrupoles. During this displacement, the DEP force becomes positive (**Erreur ! Source du renvoi introuvable**.B) and represents the crossover frequency characteristic of the cell’s intracellular composition.

By polarizing the left and right electrodes (perpendicular to the flow) with a high-frequency alternative signal, while grounding the top and bottom electrodes (parallel to the flow), we can create an effective electrical trap in the central part of the main channel, enabling the electro-manipulation of one cell at a time. For the experiment, the magnitude of the applied voltage is between 2 and 4 Vrm (root mean square voltage).

### 2.5. Statistical analysis

Statistical analyses were conducted on three independent experiments using Prism GraphPad. We employed ANOVA, Student’s t-test, and Mann–Whitney test to compare conditions, considering p-values ≤ 0.05 as statistically significant.

## 3. Results and discussion

### 3.1. Optimization of cell elution in DEP

To achieve the hyphenation between SdFFF and biosensor devices, it is necessary to align the mobile phase. For technical reasons, DEP medium was chosen as the mobile phase, as it enables the biosensor to function. However, so far, colorectal stem cell isolation by SdFFF has been carried out using PBS saline as the mobile phase [14], [15]. We therefore initially performed cell elution in SdFFF with the same field and flow rate as for PBS, using DEP as the mobile phase. As shown in Figure 1A-B, simply changing the mobile phase alters the elution profile. Under these conditions, the resolution between the dead volume containing debris and particles not retained by the field (peak no.1) and the cell peak (peak no.2) is low, making cell subpopulation sorting hazardous.

**Figure 1:**
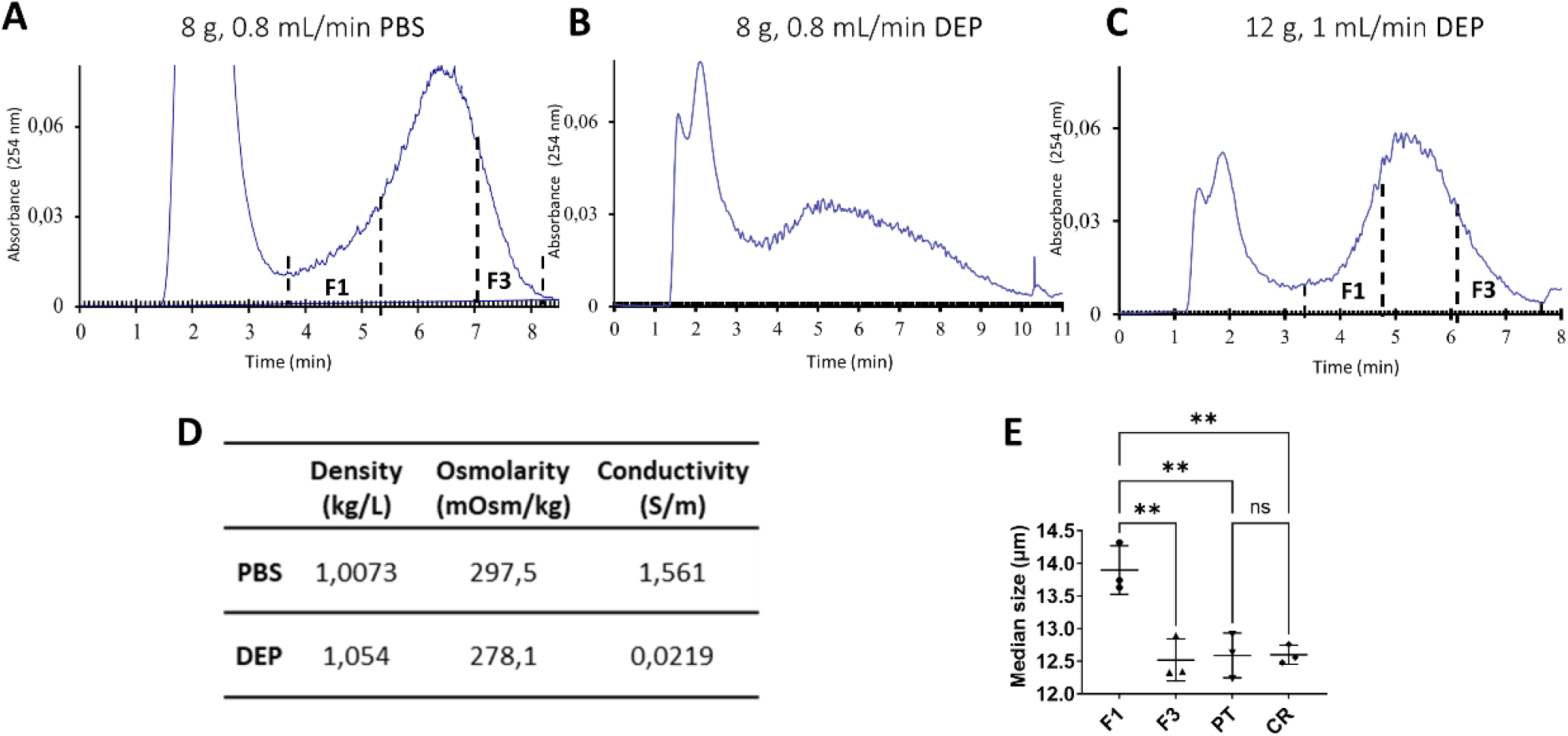
SdFFF fractograms of SW480 cell line grown in NM prior to sorting and injected at 2.5×10^6^C/mL: (A) with sterile PBS-B carrier fluid at 8g and 0.8mL/min; (B) with sterile DEP-B carrier fluid at 8g and 0.8mL/min; (C) with sterile DEP-B carrier fluid at 12g and 1mL/min. (D) Summary table of the main characteristics of DEP compared with PBS.(E) Biophysical analysis of cell size by Coulter counter of sub-population populations (F1, F3, TP, and crude). The *p* value was determined using the t Student test, ** represents p value < 0.001.

**Figure 2:**
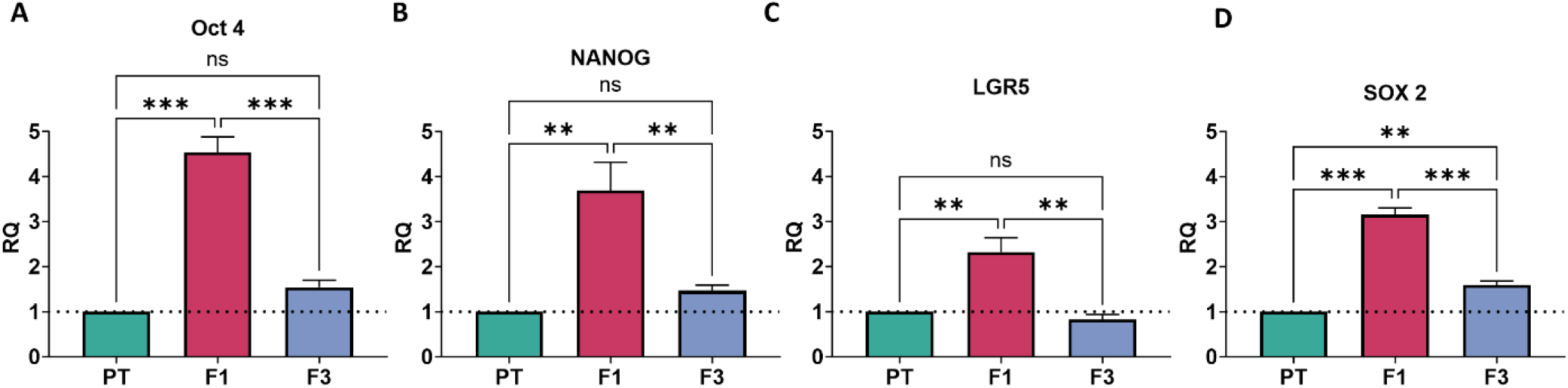
Comparative analysis of gene expression of three CSC markers: (A) Oct-4, (B) Nanog, (C) LGR5 and (D) Sox2, in the SW480 cell line, of the sub-population populations (F1, F3 and PT), measured by real-time quantitative polymerase chain reaction (PCR) and normalized compared to TP (n=4 independent experiment): The p value was determined using the t Student test. ***represents p value< 0.001 and ** represents p value < 0.001

**Figure 3:**
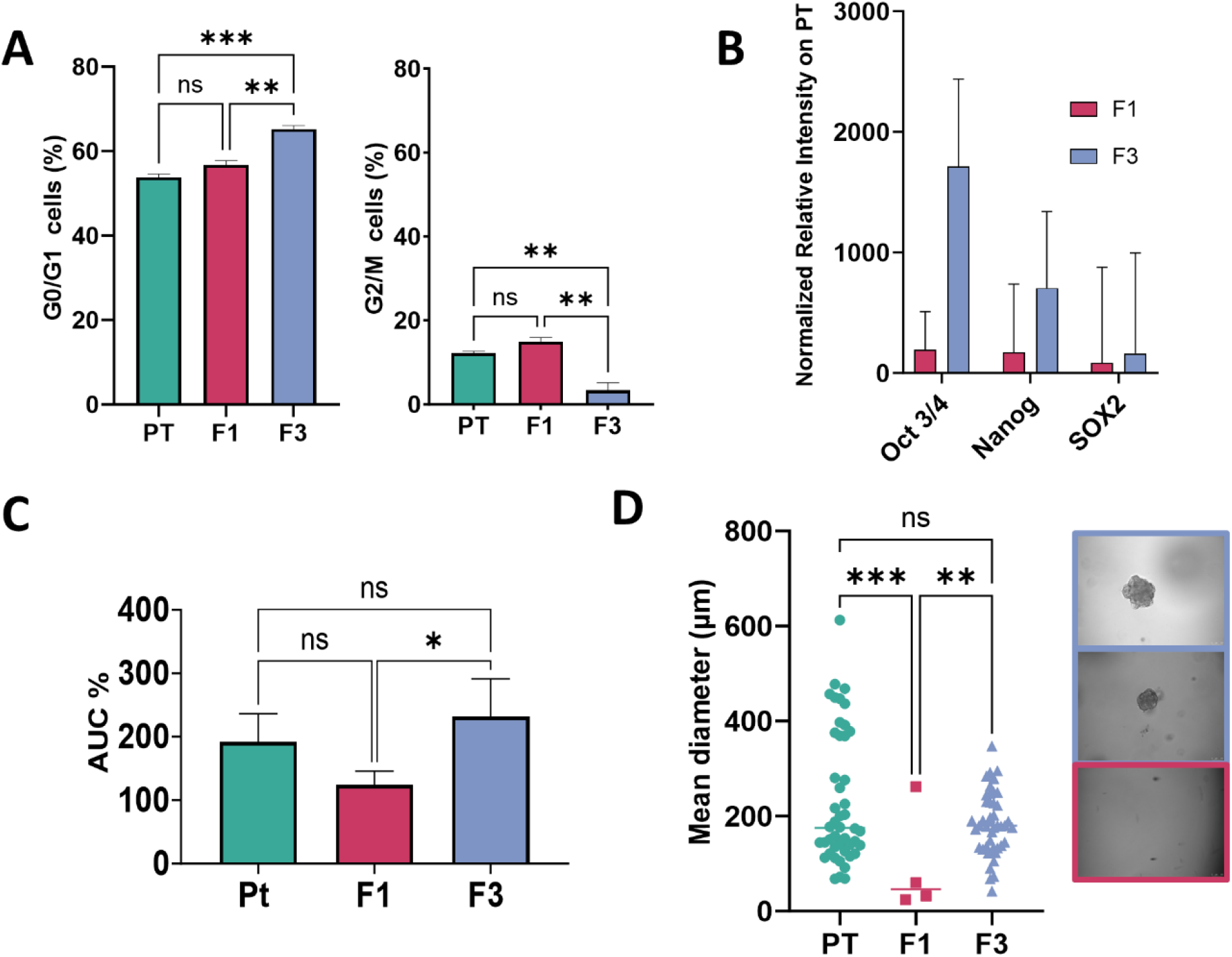
Functional characterization of the different subpopulations of SW480 sorted by SdFFF (A) Cell cycle analysis by DNA content measurement (n=3 independent experiment). (B) Proteome profiler to show the expression of a series of proteins involved in stem cell markers (n=3 independent experiment). (C) Sorted cells were seeded at 2500 cells confluence/well, and cell numbers were measured using the Incucyte live cell imaging device. Images were taken every two hours and tracked for 60 hours (SI). Experiments were carried out over 3 replicates, each in technical triplicate. The Object Count/Phase Object Count ratio was determined using Incucyte analysis. The area under the curve for each condition was then calculated and compared using a t-test. (D) Matrigel assay for colony formation between fraction at 10x magnification (n=3 independent experiment). The *p* value was determined using the *t* Student test or a two-way ANOVA. ***represents *p* value< 0.001, ** represents *p* value < 0.001 and * represents *p* value < 0.05

**Figure 4:**
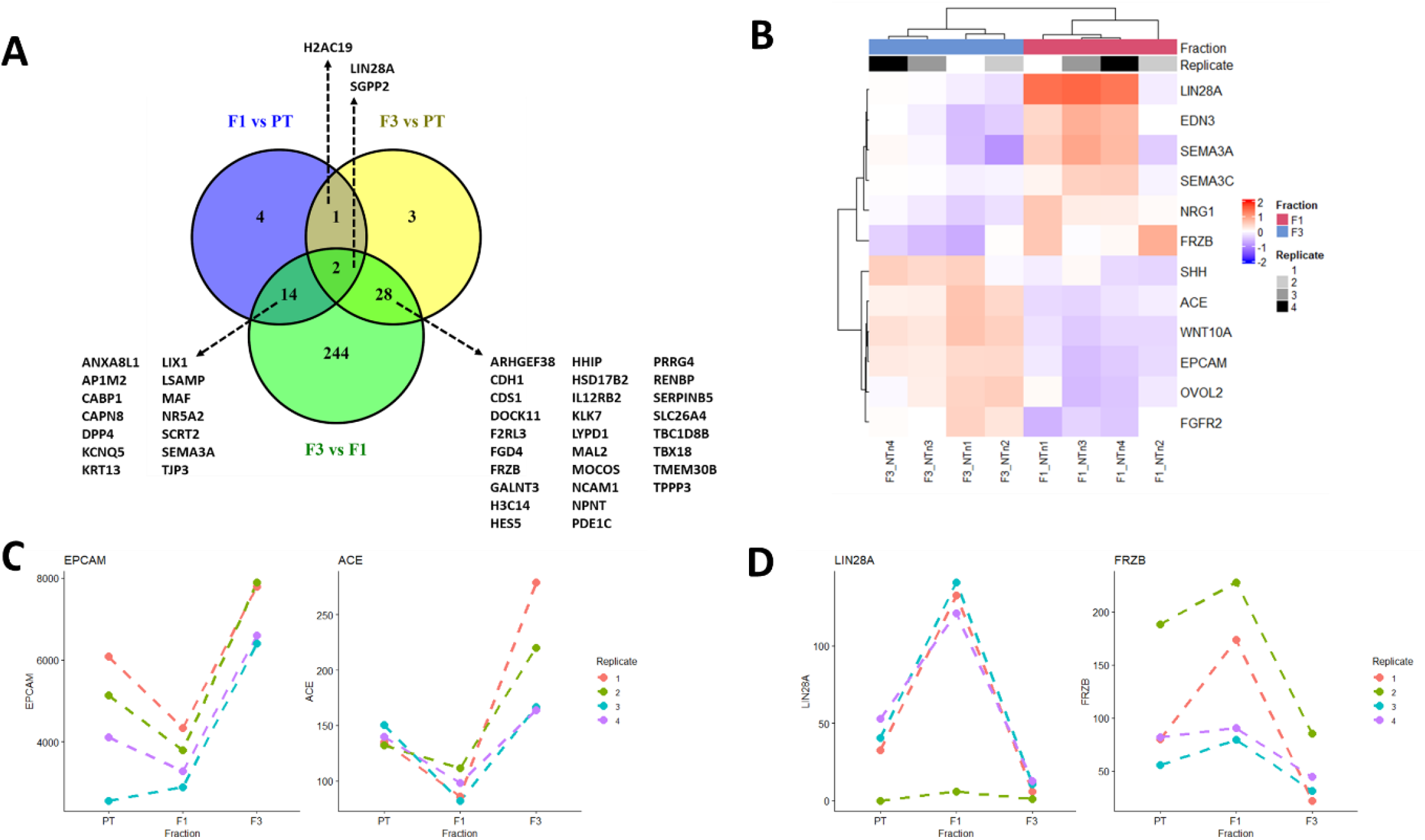
Transcriptomic analysis of sorted cells (A) Venn diagram between lists of deregulated expressed genes in pair-wise comparison of different fractions obtained after Sedimentation Field-Flow Fractionation of SW480 cell culture (n=4 independent experiment): F1 *vs* paired PT (n=21 DEGs), F3 *vs* paired PT (n=33 DEGs) and F3 *vs* paired F1 (n=288 DEGs). Gene symbol of commonly DEGs was indicated. (B) Heatmap of 12 differentially expressed genes in F3 fraction compared to paired F1 fraction, from 4 independent experiment. Gene expression were rlog transformed and median-centered and are displayed as color gradient from blue to red, reflecting low to high expression level, respectively. Genes (rows) and samples (columns) are hierarchically clustered using Pearson correlation distance and average linkage method. Samples characteristics were indicated at the top, with fraction (F1, magenta ; F3, blue) and biological replicate corresponding to 4 independent cell culture fractioned by Sedimentation Field-Flow Fractionation (1, white; 2, clear grey: 3, dark grey; 4, black). (C) Differential expression of selected genes up regulated in F3. (D) Differential expression of selected genes up regulated in F1.

**Figure 5:**
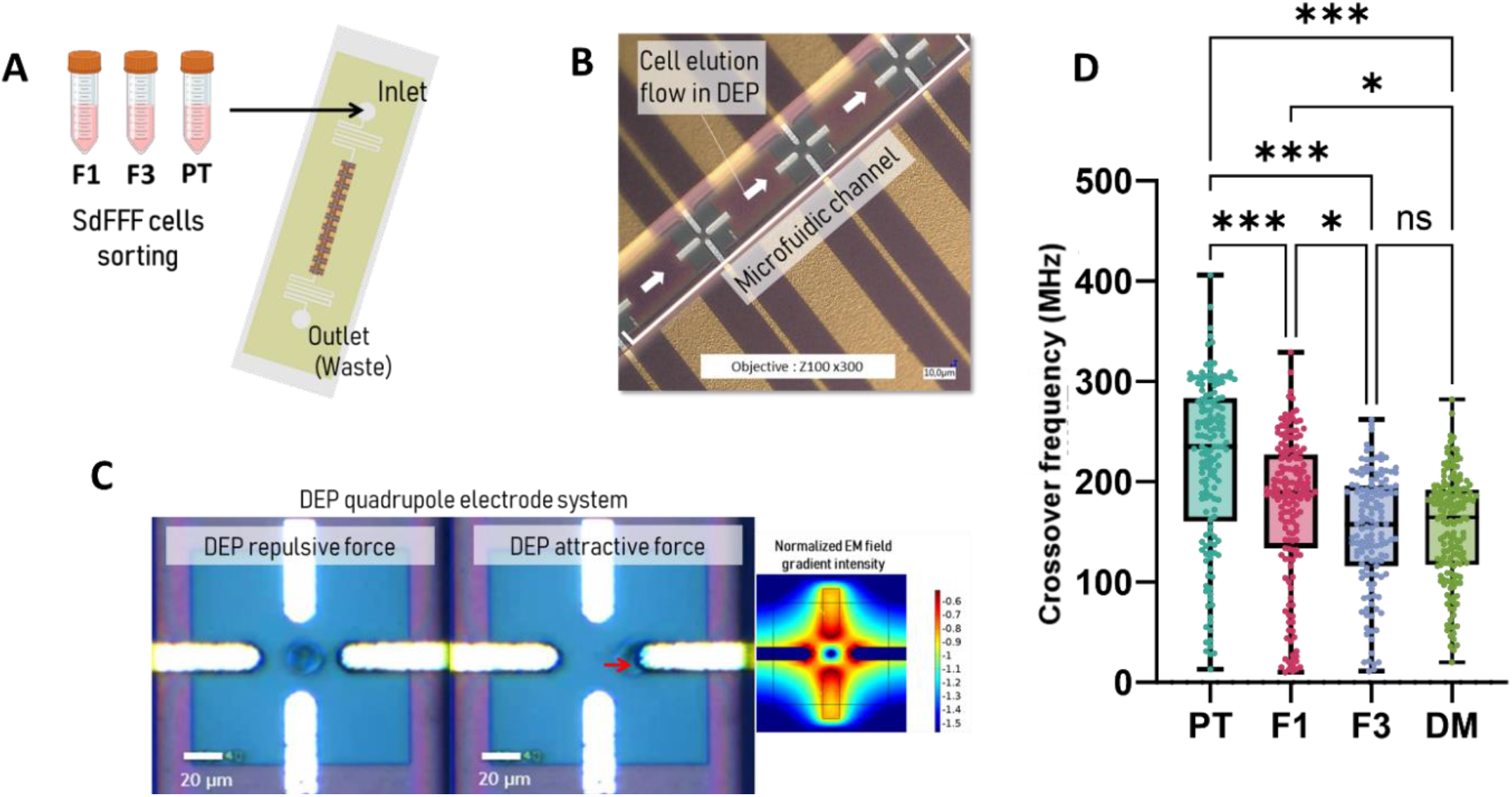
UHF-DEP Characterisation: (A) Schematic representation of post-sort cell characterization by SdFFF. (B) View of the quadrupole electrode array for DEP characterization, located at the bottom of the microfluidic channel, is visible through the PDMS system beneath the flowing cells. (C) Visualization of cellular behavior in a quadrupole electrode during UHF electromagnetic field application for DEP characterization (D) Graphic box plot representation of the HFC values of SW480 cells sorted by SdFFF and compared to culture in Defined Medium SW480 Cells. Moreover, each point represents a crossover frequency of one cell. The *p* value was determined using the one-way ANOVA test. ***represents *p* value< 0.001 and * represents *p* value < 0.05

To improve this resolution, we performed a field-flow matrix to determine the ideal elution conditions. The optimum elution conditions were found to be a field strength of 12g and a flow rate of 1 ml/min Figure 1C.

This change in conditions is explained by the variation in viscosity and density between the two solutions **Figure 1**D. Beyond elution conditions, this change in mobile phase also impacts the distribution and response of cells to the field in the separation channel. In SdFFF, a hyperlayer elution mode is applied, in which cell populations are placed according to their biophysical properties (size and density) along the flow lines, allowing larger, less dense cells to elute first, followed by smaller, denser ones (SI 3). The experimental retention rate (R_obs_) was calculated to determine the average speed and modes of elution.

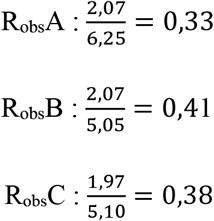

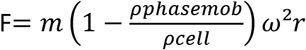, je ne retrouve pas la référence pour la masse volumique d’une cellule mais on était à 1,10g/cm^3^

This elution mode depends mainly on the biophysical properties of the cells. We collected the fractions after sorting in SdFFF and measured their mean size using a Coulter counter **Figure 1**E. As expected, the F1 fraction shows a significantly higher mean size than F3, as well as PT and CR control samples. This variation underlines the effectiveness of SdFFF for size-based sorting. Furthermore, the low size observed for the PT and CR controls can be explained by the time spent in the DEP medium, which appears to decrease cell size over time (SI 2).

Although this information shows the impact of the new mobile phase on cell elution in SdFFF, it is essential to validate that this change does not alter the efficiency of SdFFF for CSC sorting. To achieve this, a number of biological characterization methods need to be implemented, including phenotypic, functional and, for the first time, transcriptomic assays.

### 3.2. Phenotypic and functional characterization of sorted cells

In solid tumors such as colorectal cancer, a great diversity of cells is present, including cancer stem cells (CSCs), whose plasticity contributes to tumor heterogeneity. These properties make their characterization complex, requiring the use of CSC-specific phenotypic markers and functional assays to better target analyses.

To phenotypically identify CSCs in colorectal cancer, several markers, including the transcription factors Sox2, Nanog and Oct4, as well as specific genes, are used. We conducted RT-qPCR analysis of mRNA expression levels of these CSCs in populations isolated by SdFFF in the SW480 line **Erreur ! Source du renvoi introuvable**.. The overexpression of these markers in the F1 fraction, after normalization against the total population and comparison with F3, indicates a potential enrichment of CSCs in F1. This results confirmed the previous tendency published by Hervieux et al [15] obtained with PBS as mobile phase, demonstrating the great interest in DEP using.

Nevertheless as also described [3], [5], the CSC characterization is based on functional properties, including their proliferative activity, the expression of specific proteins and their clonogenic capacity. For example, CSCs have the ability to remain in a quiescent state in the tumor niche, which favors their multipotency within the tumor. In general, these cells are predominantly in the G0/G1 phase of the cell cycle, while differentiated and proliferative cells are in the G2/M phase. In our study, cell cycle analysis by flow cytometry shows that 65% of cells in the F3 fraction are in the G0/G1 phase, with less than 5% in the G2/M phase, while no significant difference is observed in F1 compared to the total population **Erreur ! Source du renvoi introuvable**.A. These results suggest that the F3 fraction isolated by SdFFF is enriched in CSCs in G0/G1 phase, and therefore exhibits quiescence capacity.

To check for the presence of proteins characteristic of CSCs such as Nanog, Sox2 or Oct4, we carried out a proteome profiler analysis. Analysis of the relative intensities of dot-blots showed a general orientation towards the expression of these proteins, mainly in the F3 fraction **Erreur ! Source du renvoi introuvable**.B.

CSCs are also known for their high proliferative capacity. When observing cell proliferation by IncuCyte, we found that the F3 fraction had a higher proliferative capacity than the other fractions **Erreur ! Source du renvoi introuvable**.C.

Finally, one of the reference criteria for identifying CSCs is their clonogenic capacity, reflecting their potential for self-renewal. To this end, we cultured the sorted cells in a Matrigel matrix. Our solid-state assay revealed that the F3 and PT subpopulations are the most capable of initiating tumorosphere formation **Erreur ! Source du renvoi introuvable**.D.

This series of biological characterizations shows that after culturing SW480 cell lines in normal medium (NM), the F3 fraction is enriched in CSCs, while the F1 fraction appears to be enriched in progenitor stem cells, as indicated by its high mRNA level. Consequently, SdFFF is validated as a label-free cell sorting method, using DEP-B as carrier liquid instead of PBS.

In order to deeply characterize cell subpopulation, we performed for the first time a RNA Seq analysis.

### 3.1. Transcriptomic characterization

The use of these biological tests to characterize CSCs is commonplace, but relies on the assumption that they actually identify CSCs. We performed 4 replicates of the sub-population collection to analyze their transcriptomic content. As each fraction in each replicate is derived from the same sample, we were able to perform a paired analysis (SI 5). This approach, well suited to the small number of replicates and to inter-individual variability, enables us to better control the effects linked to differences between individuals, and thus reinforces the robustness of the results obtained. For this analysis, a total of 296 genes were studied, 288 of which were differentially expressed between F1 and F3 (**Erreur ! Source du renvoi introuvable**.. A) (SI 6). Of these, 141 were up-regulated in F3 and 147 down-regulated in F1. Enrichment analysis revealed 12 genes involved in CSC differentiation (SI 7). Half of these genes are over-expressed in F1 and under-expressed in F3, as shown in **Erreur ! Source du renvoi introuvable**.. B.

These two groups of strain-related enrichment exhibit distinct functions associated with the fractions in which they are enriched. In Fraction 3, genes such as SHH, WNT10A, ACE, OVOL2, EPCAM, and FGFR2 play a crucial role in maintaining the strain traits related to cancer stem cells (CSCs). Specifically, SHH and WNT10A activate the Hedgehog and Wnt signaling pathways, which are essential for CSC proliferation and maintenance. Additionally, EPCAM supports CSC adhesion and self-renewal, while also playing a role in epithelial-mesenchymal transition (EMT), a process that enhances tumor progression.

In contrast, genes enriched in Fraction 1 (LIN28A, SEMA3C, EDN3, FRZB, SEMA3A, NRG1) are primarily involved in early cellular processes and the initiation of stem cell traits. For instance, LIN28A is known for its role in embryogenesis and cellular reprogramming, highlighting its function in stemness maintenance [33], [34]. This difference between F1 and F3 suggests that F1 genes contribute to the foundational properties of CSCs, whereas F3 genes are more directly involved in supporting their persistence and proliferation in the tumor microenvironment. These transcriptomic results confirm what was observed during the biological, phenotypic and functional characterizations, while suggesting that markers such as EpCam could be used as discriminating markers of SdFFF sorting efficiency.

Biological characterizations reveal that after culture of the SW480 line in normal medium (NM), the F3 fraction shows enrichment in CSCs, while F1 shows characteristics of progenitor stem cells, notably by its high mRNA expression. These results validate the efficacy of the SdFFF label-free sorting method, using DEP-B as the mobile phase instead of conventional PBS, for isolating colorectal cancer CSCs.

### 3.2. Characterization of UHF-DEP biosensor

Now that SdFFF has demonstrated its effectiveness in isolating CSCs from the SW480 line, using DEP as the mobile phase, we can turn our attention to the coupling of SdFFF and UHF-DEP. To this end, the F1, F3 and PT sub-populations are characterized by the UHF-DEP by measuring the value at which the cell held at the center of the quadrupoles changes from a repulsed state to a state attracted to the electrode, corresponding to the high-frequency crossing (HFC).

Preliminary testing of the PT coupling revealed a critical point in the protocol not observed in Glioblastoma studies[13], [19]. For SW480 cells, it was necessary to recover them from the culture medium during the collection of subpopulations at the exit of SdFFF. They were then centrifuged and resuspended in DEP medium prior to biosensor characterization (SI 4). Without this step, the cells showed aberrant HFCs, due to their fragile state. We therefore optimized this procedure, which was not necessary in the first place, to ensure that measurements would not be biased, while retaining the DEP-B mobile phase for both techniques.

As a first step, SW480 subpopulations sorted by SdFFF and grown in NM medium were characterized via the UHF-DEP biosensor. As shown in **Erreur ! Source du renvoi introuvable**.. **Erreur ! Source du renvoi introuvable**.D, median HFC values differed between each subpopulation: populations F1 and F3 both had significantly lower medians (190 and 157 MHz respectively) compared with the PT, located at 237 MHz, but also with each other. To complete this analysis, we compared these subpopulations with a population of cells grown in a defined medium, known to enrich the population in CSC [35] and considered as a gold standard. By measuring the DM condition in HFC, we were able to pinpoint the crossover point for CSCs. As shown in **Erreur ! Source du renvoi introuvable**.. D**Erreur ! Source du renvoi introuvable**., there is no significant difference between cells grown in defined medium and those of fraction F3, while a significant difference is observed between cells in defined medium and those of fraction F1. These significant differences between F1 and PT and between PT and F3 highlight the presence of two populations close to the HFC of CSC-enriched cells, represented by the defined medium condition. These results also confirm the observations made during biological characterization, namely that the SW480 sub-population sorted by SdFFF in DEP corresponds to a CSC population, while the F1 fraction seems to approach it as a progenitor cell population. These results also confirm the observations made during biological characterization: the SW480 subpopulation sorted by SdFFF and UHF-DEP corresponds to a population of CSCs, while the F1 fraction seems to approximate a population of progenitor cells. We can therefore conclude that the HFCs obtained are specific to each subpopulation isolated by SdFFF. Moreover, these HFCs can be associated with the results of the RNAseq analysis. Indeed, in the enrichment analysis (figure X SI), we observe that the pathways of ion channel activation, receptor signaling, cytokine binding, cell surface genes and calcium signaling are particularly activated, playing a key role in cancer development. Examining the relationship between CSC and UHF-DEP biosensor function, we find that, although HCF is representative of the cells’ internal composition, it is possible that the activity of channels present on the cell membrane surface contributes to the observed decrease in HFC in the F3 population. Studies have also shown that Ca^2+^, Mg^2+^, K^+^ and Zn^2+^ ion channels play a crucial role in the maintenance and survival of CSCs [36], notably by supporting their capacity for self-renewal. These membrane transporters are also known to influence the bioelectrical properties of cells [37], aspects that can be detected and analyzed by the UHF-DEP biosensor.

## 4. Conclusion

Cancer stem cells (CSCs) play a crucial role in the development of colorectal cancer recurrences. However, current isolation methods, such as flow cytometry (FACS/MACS), as well as biological characterization techniques through phenotypic and functional tests, do not allow for rapid and complete identification of CSCs, while risking modifications to these cells. Previous studies have demonstrated the effectiveness of SdFFF and UHF-DEP biosensors for isolating and characterizing CSCs in the context of glioblastoma [13]. We aimed to adapt this work to colorectal cancer to validate the universality of this coupling method.

Although these techniques have already proven their effectiveness, the standardization of the mobile phase necessary for their coupling required new biological calibration. Results showed that, despite the changes introduced by this technical modification, SdFFF remains an effective tool for discriminating a population of precursor CSCs in F1 and a population of functional CSCs in F3 as see in previous study [15].

Moreover, characterization of the subpopulations F1, F3, and PT by UHF-DEP revealed a significant decrease in HCFs in the fractions of interest (F1 and F3), which can be associated with the reference condition in a defined medium, thereby reinforcing the relevance of the UHF-DEP biosensor for discriminating CSCs. Transcriptomic analysis of the subpopulations also confirmed the biological and physical observations made, highlighting a differential enrichment of differentiation genes between F1 and F3. This indicates that F1 retains tumorigenic and tumor-initiating capabilities, while F3 exhibits proliferative, differentiative, and expansion abilities, as evidenced by the overexpression of EpCAM [38].

Finally, in addition to biological validation, conducting a transcriptomic analysis has, for the first time, established a link between the decrease in HCFs and CSCs. We identified an overexpression of pathways involved in the modulation of membrane channels characteristic of CSCs [37], opening multiple options for better understanding the intracellular changes identified by UHF-DEP.

## Supporting information

Supplementary datas

