## Supplementary datas for "Towards a standardised method for the characterisation and isolation of colorectal cancer stem cells by SdFFF and UHF-DEP: highlighted by transcriptomic analysis"

**Supplementary figure legends:**

**Supplementary figure 1.**

Experimental diagram of the article

**Supplementary figure 2.**

Evolution of cell size as a function of time in contact with DEP or PBS

**Supplementary figure 3.**

Graph of viscosity and stress as a function of shear rate

**Supplementary figure 4.**

Optimization of SdFFF output cell characterization for cell detection by UHF-DEP

**Supplementary figure 5.**

Principal component analysis of transcriptomic data obtained from 4 independent SW480 culture, fractioned in PT, F1 and F3 fractions by Sedimentation Field-Flow Fractionation. Shape of symbol corresponds to biological replicate and color of symbol correspond to fraction. Normalized counts with rlog transformation were used for PCA analysis. The variability between biological replicates prevents the fractions from being properly separated, which has led us to carry out a differential analysis in paired condition in the rest of our analyses

**Supplementary figure 6.**

Enrichment analysis of Gene Ontology Biological Process (BP), Molecular Function (MF) and Cellular Component (CC) and associated to pathway database (Reactome and KEGG). Enrichment analysis was performed from 288 DEGs in F3 *vs* F1 on Database for Annotation, Visualization and Integrated Discovery (DAVID) online tool (https://david.ncifcrf.gov/) with cluster analysis of enriched terms performing to recover redundant terms. The selected term corresponds to the one with the most significant p-adjusted within each top-6 enrichment cluster

**Supplementary figure 7.**

Visualisation of gene expression level of 15 genes by graphical plot (y axis = normalized counts)

**Supplementary figure 1.**

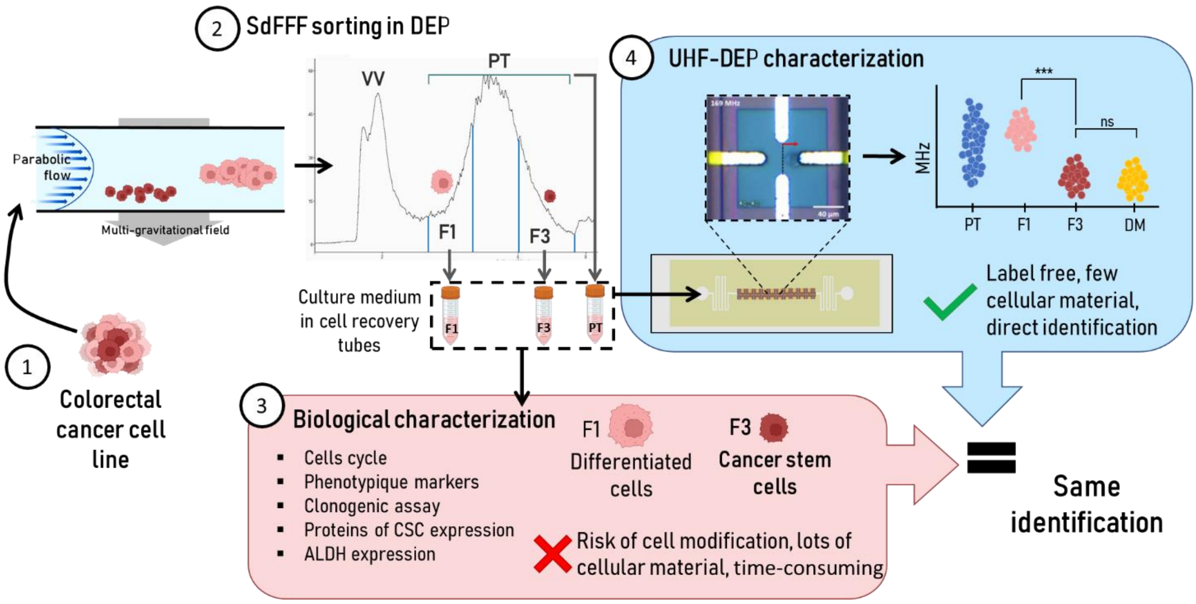


**Supplementary figure 2.**

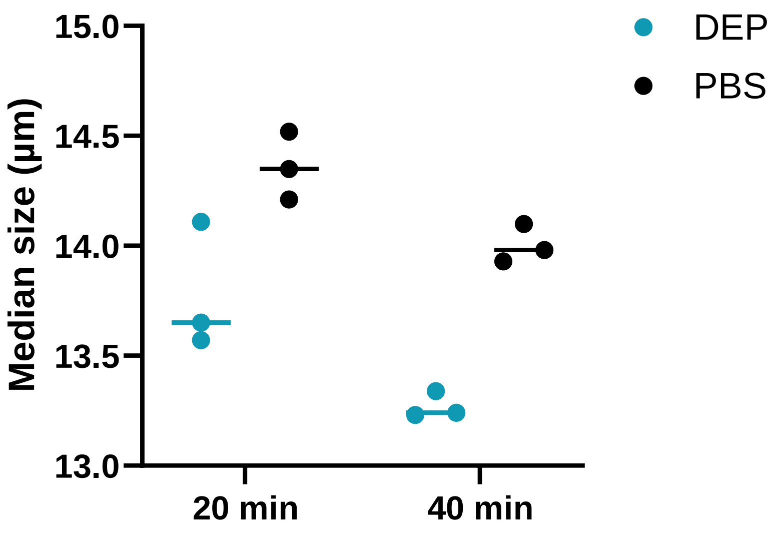

**Supplementary figure 3.**

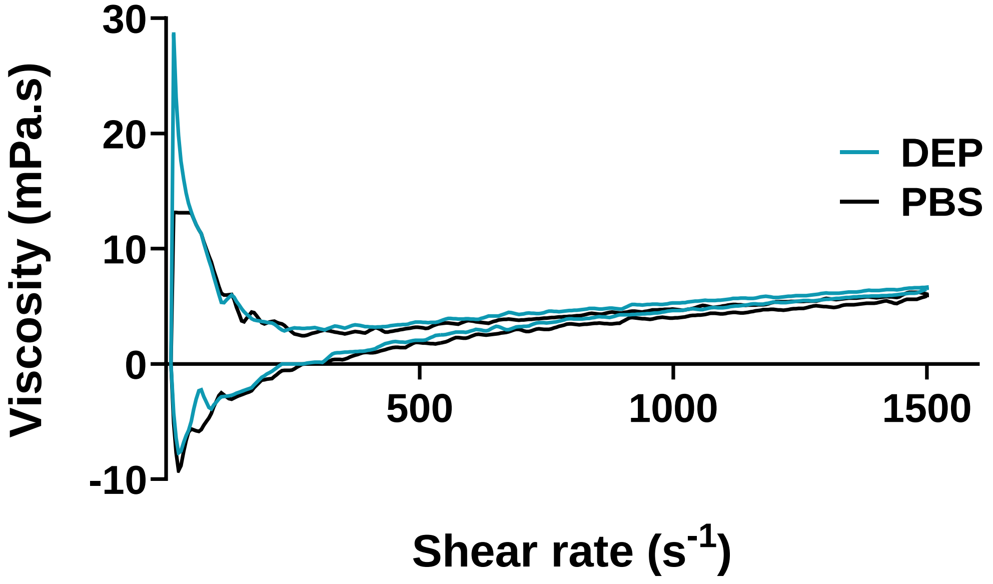


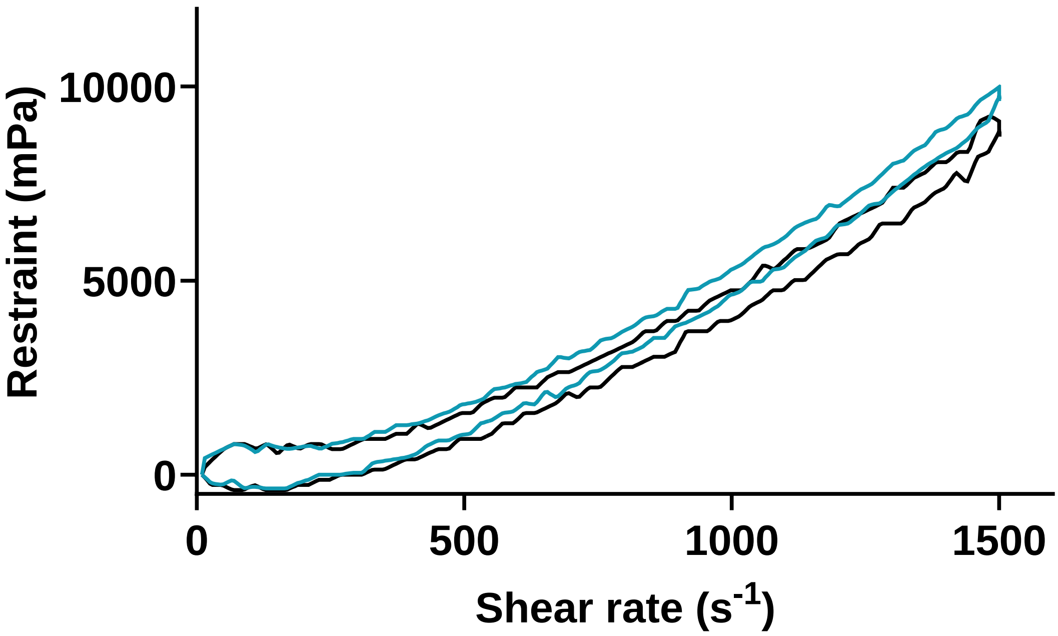

**Supplementary figure 4.**

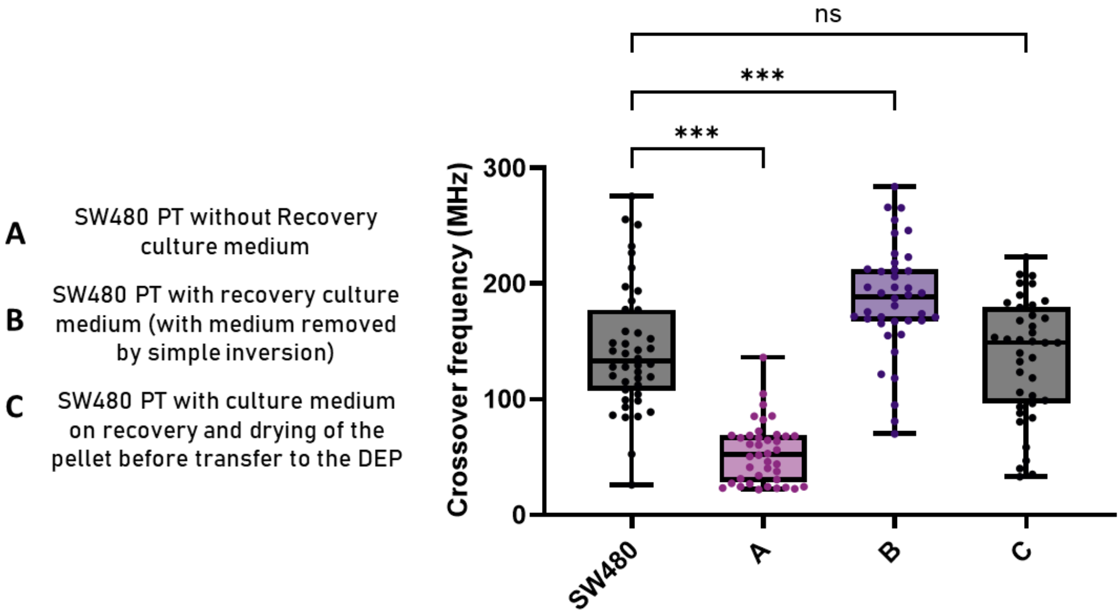


<u>Optimization of SdFFF output cell characterization for cell detection by UHF- DEP</u>

**Supplementary figure 5.**

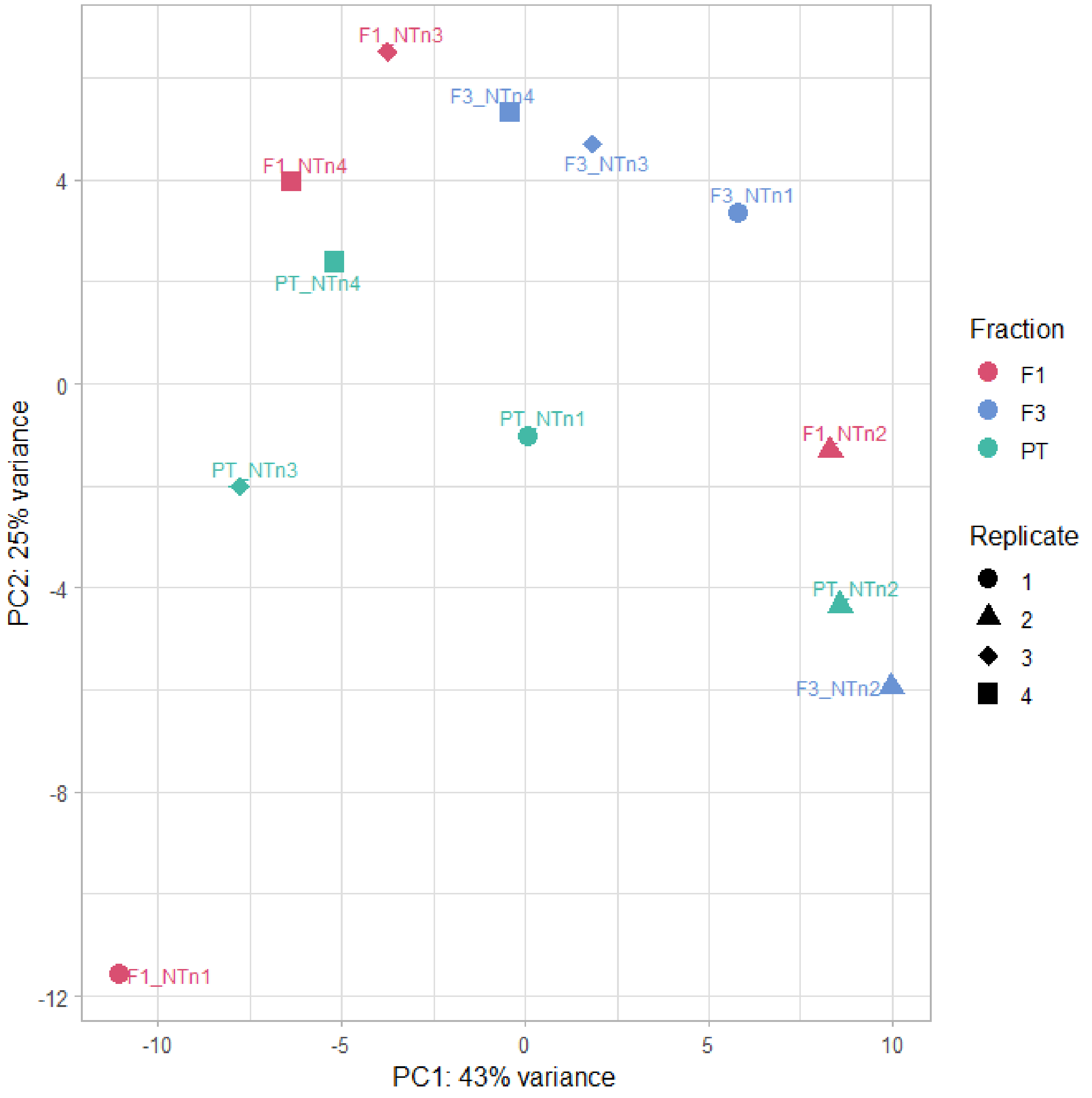

**Supplementary figure 6.**

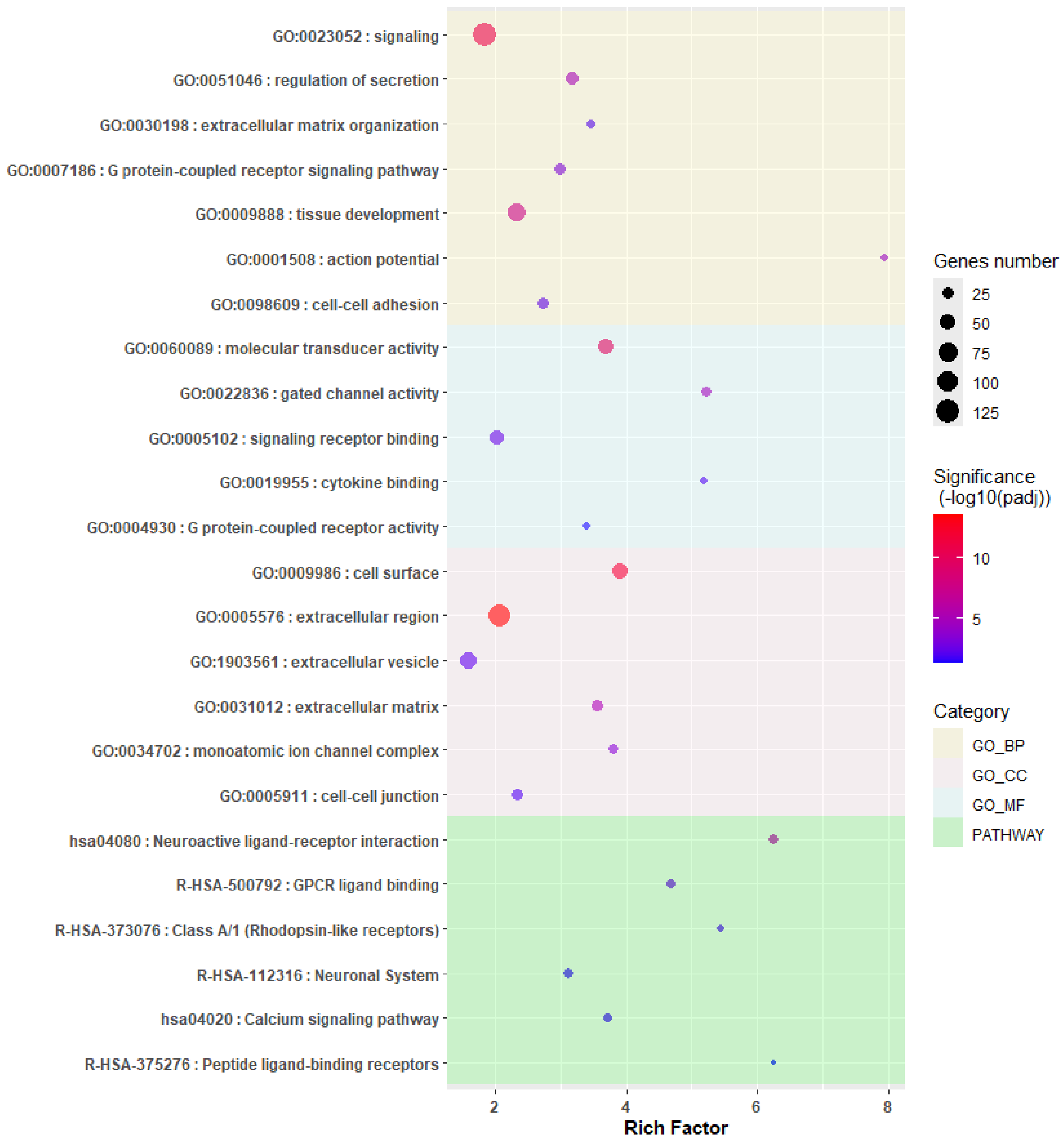

**Supplementary figure 7.**

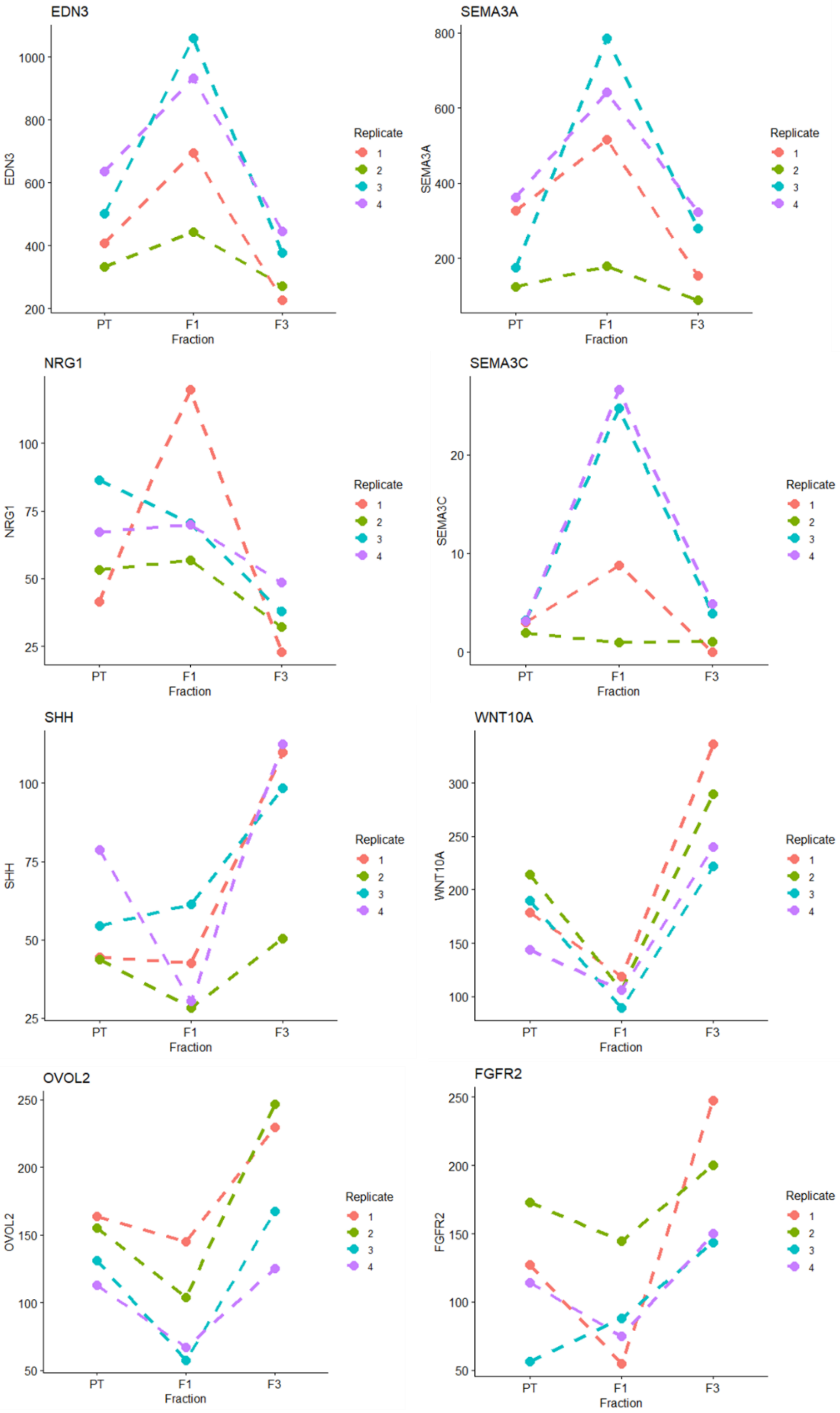

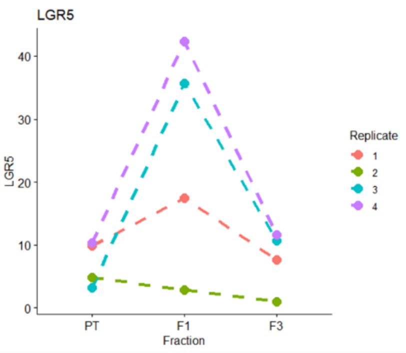


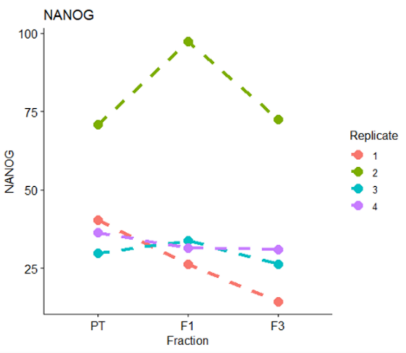


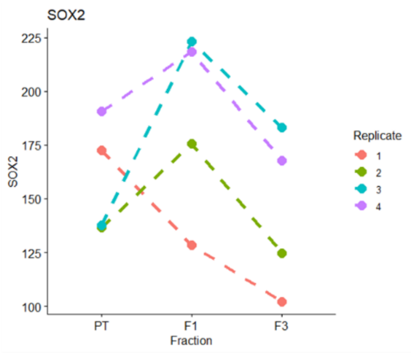


**LGR5**
log2FC = -1,596092698
Padj= 0,03865261

**NANOG**
log2FC = -0,385099072
Padj = 0,58905252

**SOX2**
log2FC = -0,373090327
Padj = 0,31706570
